# The Nucleoporin MLP1 of the nuclear basket is essential for nuclear integrity and ploidy maintenance in *Trypanosoma brucei*

**DOI:** 10.64898/2026.01.09.698594

**Authors:** Akila Yagoubat, Lucien Crobu, Slavica Stanojcic, Nada Kuk, Amélie Sarrazin, Marie-Pierre Blanchard, Patrick Bastien, Maude F Lévêque, Laurence Berry, Yvon Sterkers

**Author notes:** Université Paris Cité, CNRS, Institut Jacques Monod, Paris, France.

## Abstract

*Trypanosoma brucei* is a divergent eukaryote parasite responsible for neglected tropical diseases in human and animals, sleeping sickness or human African trypanosomiasis and nagana respectively. Besides scientific interest, understanding the specific features of its biology has medical and economical relevance. Nuclear pore complexes (NPCs) are large multiprotein channels embedded in the nuclear envelope that regulate nucleo-cytoplasmic transport. Beyond this role, NPCs also participate in essential nuclear processes such as chromosome segregation, transcription, and cytokinesis. Here, we showed that Myosin-like protein-1 (MLP1) localized to the nuclear basket of NPCs in *T. brucei*. Silencing of TbMLP1 by RNA interference in *T. brucei* procyclic cells caused marked defects in parasite growth, severe impairment of messenger RNA export, disorganization of nuclear structure, and pronounced genomic instability. Flow cytometry and FISH analyses revealed abnormal DNA content and a sharp reduction in disomic cells, accompanied by increased numbers of monosomic, trisomic, and polysomic cells, indicating an untolerated aneuploidy.

These findings reveal that TbMLP1 links NPC function to multiple key cellular pathways. We provide new insights into the mechanisms that preserve nuclear architecture, nuclear envelope morphology, genome stability, and faithful chromosome segregation, including effects on kinetochore distribution and organization of the mitotic spindle.

**Author Summary:** *Trypanosoma brucei* is the microscopic parasite responsible for sleeping sickness in humans and nagana in cattle; learning more about its basic biology may help identify weak points that could one day be targeted in new treatments. In this work, we studied a protein called MLP1, which sits at the nuclear pores—the gateways that control what enters and leaves the nucleus. Our findings revealed that MLP1 is essential for ensuring proper movement of messenger RNA, keeping the nucleus stable, and allowing chromosomes to be shared equally when cells divided. In total, we gained new insight into the basic biology of a parasite that remains a major health concern in many regions of the world, showing how a single structural protein can influence many vital processes within it.

## Introduction

*Trypanosoma brucei* represents an unconventional eukaryotic model for biological studies. This protozoan parasite causing Human African Trypanosomiasis (HAT or sleeping sickness) and Animal African Trypanosomiasis (AAT or Nagana in animals) have an important global burden on health and economic progress. *T. brucei* is a divergent unicellular eukaryote, far from other model organisms (1). This divergent eukaryote undergoes a closed mitosis during which the nuclear envelop remains intact and the mitotic spindle is assembled intranuclearly like in yeast (2, 3). It also shows very original and unusual features. *Trypanosoma* has two independent but coordinated cell cycles (nuclear and mitochondrial) (4). The genome is organized in large unidirectional polycistronic units that are composed of tens or hundreds of functionally unrelated genes (5, 6). Transcription is RNA polymerase II directed whereas theres is a near absence of promoters; gene expression regulation occurs mainly at the posttranscriptional level (7). Nuclear pore complexes (NPCs) are macromolecular structures that perforate the nuclear envelope (NE) forming the unique sites of exchange between the cytoplasm and the nucleus. NPC is one of the largest protein complexe in eukaryotes; it is constituted by ∼30 different proteins (in yeast and vertebrates). These structural protein units are called nucleoporins (NUPs) (8, 9). The detailed structure and folding of NPCs have been well established in *Saccharomyces cerevisiae* (10) and in mammals (11, 12). The NPC structure consists of a symmetrical part known as the core complex anchored in the nuclear envelope (8) and an asymmetrical part which is composed of filaments emerging from the cytoplasmic and nuclear sides. While the cytoplasmic filaments emerge freely from the NPC, the nuclear filaments are attached to the NE in the nuclear side to form the nuclear pore basket (13). The general structure/folding of the NPCs is well conserved between yeast, vertebrates and plants (14) and even in more divergent eukaryotes like *T. brucei* (15). Besides their roles in nucleo-cytoplasmic transport, there are increasing evidences showing that NUPs have a pivotal role in maintaining nuclear organization and genome integrity (reviewed in (9)). For example, the spatial organization of other nuclear complexes such as the transcription export complex (TREX) (16), DNA repair (17, 18), cell cycle progression, chromosome segregation during mitosis (19–21) and gene transcription (as reviewed in (22, 23)). One of the most important parts of the NPCs is the nuclear basket. The nuclear basket plays a key role as a multifunctional platform for various nuclear processes (21, 24). It contributes to the (i) maintenance of the nuclear organization/stability (25), (ii) control telomere length and repair of DNA damaging (26–30), (iii) spatial regulation of the Spindle Assembly Checkpoint (SAC) (31–34). Sequence homology search failed to identify any clear NUPs homologs in trypanosomatids genome indicative of a highly sequence divergence. However, in 2009, 22 TbNUPs were identified using proteomic approaches followed by *in situ* tagging and localization in *T. brucei* (15). Despite the low primary sequence similarities, TbNUPs share secondary structural organization with other model organism’s NUPs (15, 35) with the exception of the peripheral regions such as the nuclear basket which is more divergent (36–38). The nuclear basket is mainly formed by TPR (translocated promoter region) proteins encoded by multicopy genes in mammals. TPR homologs in fungi and trypanosomatids are named myosin-like proteins (MLP1 and MLP2). There is clear evidence that TbNUP92/TbMLP2 is implicated in chromosomal distribution and genome stability (37, 39). MLP1 “myosin like protein in Trypanosomatids seems to be specific to kinetoplastid at both genomic sequence and domain organization levels. In this work we have characterized NUP110/MLP1 in *T. brucei.* Our results showed that TbMLP1 is localized to the nuclear pore basket all along the cell cycle; that TbMLP1 is essential for cell survival and correct cell cycle progression in *T. brucei*. We demonstrated that TbMLP1 is required to maintain nuclear architecture integrity, the NE morphology and the ploidy. Further, TbMLP1 appeared to be important for chromosomes segregation process *via* a defect in kinetochore distribution and mitotic spindle’s network organization.

## Methods

### Parasites in vitro culture

Procyclic forms of the *T. brucei* Lister 427 wild type were used as control strain and *T. brucei* Lister 427 29-13 co-expressing the T7 RNA polymerase and the tetracycline repressor was used to generate the TbMLP1 RNAi knockdown cell lines. To generate the strain co expressing the T7TR system and the Cas9, Tb Lister 427 29-13 (T7TR) was transfected with an episome coding for the SpCas9-HA and the puromycin resistance cassette (40). The Tb427-T7TR-Cas9 was then used to generate RNAi and tagged cell lines. All cell lines were grown at 27°C in SDM-79 supplemented with 10% heat inactivated tetracycline free FBS, 7 µg.mL^-1^ hemin, 30 µg.mL^-1^ of hygromycin (InvivoGen) and 10 µg.mL^-1^ of geneticin (InvivoGen) for the Tb Lister 427 29-13 (T7TR) line in addition to 30 µg.mL^-1^ of puromycin (InvivoGen) for the strain expressing Cas9. To establish cell growth curves, every second day, cells were counted using a Denovix CellDrop 2 Channel Fluorescence and BrightField and diluted to 3×10^6^ cells/mL. In *T. brucei*, cell cycle phases can be followed by the number and position of DNA-containing organelles, *i.e.* nuclei (N) and kinetoplasts (K). This feature is used to determine the Nucleus/Kinetoplast pattern (NK pattern) after staining cells with DAPI. The NK pattern corresponds to the proportions of 1N1K, 1N2K, and 2N2K cells (normal events) and abnormal cells containing >2 K and/or >2 N in the population. More than 200 cells were counted for each time point.

### MLP1 RNAi and tagging strains in T. brucei

All sequence analysis were performed using TriTrypDB (41, 42), the integrated functional genomics resource for kinetoplastida; part of vEuPathDB, the eukaryotic pathogen vector and host bioinformatics resource center (43).

#### MLP1 RNAi in *T. brucei*

The RNAi targeted sequence of MLP1 in *T. brucei* 427 (TbMLP1) were identified as described in ((44) and https://dag.compbio.dundee.ac.uk/RNAit/) and PCR- amplified from genomic DNA of *T. brucei* Lister 427 wild type cell line. For TbMlp1, the oligonucleotides primers used were 5’ GGGCCGCGGGGGAAGGTGCAATACAGGAA 3’ and 5’ GGGAAGCTTGTTGCGATGAAAACAAAGCA 3’. The PCR product was cloned into pGEM-Teasy (Promega). The target sequence from TbMLP1 was then subcloned into p2T7tiB vector using the restriction sites HindIII and SacII to generate TbMLP1 p2T7 RNAi vector. Prior to transfection, 10 mg of TbMLP1 p2T7 vector were linearized using NotI and purified using the Kit Wizard®SV Gel and PCR Clean-Up System (Promega®) following the productor instructions and heat-sterilized at 75°C for 10 min. Parasite transfections were performed as described previously (45), using the Amaxa Nucleofector 2b and X-001 programs. A total of 2.10^7^ cells from exponential cell culture of the *T. brucei* Lister 427 29–13 (T7TR) cell line was used per transfection. Transfected cells were left to recover for 18-24h then selective pressure was added (5 µg.mL^-1^ of phleomycin). Induction of the RNAi cell lines was performed by addition of 3 µg.mL^-1^ of tetracycline.

#### *In situ* tagging in *T. brucei*

*In situ* tagging in *T. brucei* wild type strains (without Cas9) was performed using the long primers PCR tagging strategy and the pMOTag4G vector as a template (46). Primers used for *in situ* tagging of TbMLP1 were 5’ATGCGACTACTGCACGTCAACAAGCAACTTGTGGAGAGAGTCAAAACCAGTCGAACTGAAGGAGAATCCCAGTCCAGTGGTACCGGGCCCCCCCTCGAG3’ and 5’TACACGAATTGTCATACAACCTGACTAGCAGACGTAAGGCGCTACGAACCTTTACTGTGGTTCAAACAAAAATGGCGGCCGCTCTAGAACTAGTGGAT 3’. The primers contained homology regions of 80 bp. The linear PCR products containing the two-homology regions, the tag and a drug selection marker were amplified, purified and sterilized by ethanol precipitation. *T. brucei* Lister 427 procyclic stage cells were transfected by electroporation with 10µg of PCR product and allowed to recover. Protein localization was verified by immunofluorescence assay (IFA).

#### Gene tagging using CRISPR-Cas9 in Tb427-T7TR-Cas9

For gene tagging coupled with the RNAi knockdown of MLP1 the cell line Tb427-T7TR-Cas9 was used. The TbMLP1 RNAi was generated as explained above and the tags were introduced using the PCR-based CRISPR-Cas9 strategy for which the sgRNA and the donor DNAs containing the tag and the selection marker were amplified and transfected as PCR products (Beneke et al., 2017).

#### Gene tagging in TbMLP1 RNAi

The localization of the following proteins were analyzed MLP1 (Tb427tmp.03.0810), TbNUP109 (Tb427tmp.01.7630), TbNUP98 (Tb427.03.3180), TbNUP-1(Tb427.02.4230) (36), KINF (Tb427.03.2020) (47, 48), KKT2 (Tb427tmp.01.2290) (49). All the primers for sgRNA and donor DNA were designed through the LeishGEdit website (www.leishgedit.net) (45). The primers sequences are available in the S1 Table). 20 ng of circular pPoT vectors (pPOT-NeonG phléomycin and pPOT-HA hygromycin) were used as templates to amplify the donor DNAs for gene tagging. For PCR product amplification, 0.2 mM dNTPs, 2 µM each of gene-specific forward and reverse primers and 1-unit Phusion Polymerase (NEB) were mixed in 1 X Phusion reaction buffer, 50 µL total volume. PCR steps were 30 s at 98 °C followed by 35 cycles of 30 s at 98 °C, 30 s at 61 °C, 1min at 72 °C followed by a final elongation step for 10 min at 72 °C. Except the plasmid templates, similar PCR mix was used to amplify the sgRNAs. Final PCR products of donor DNAs and sgRNAs were purified using the Kit Wizard®SV Gel and PCR Clean-Up System (Promega®) and heat-sterilized at 75 °C for 10 min. Five µg of each donor DNA and 5 µg of the specific sgRNA were used for each transfection. Transfections were performed in the Amaxa Nucleofector using the X-001 program as described above and in (45). After recovery tag integration was checked using PCR and immunofluorescence localization. Protein behaviors and localization were followed in non-induced or tetracycline induced TbMLP1 RNAi cells.

### qRT PCR

To control the efficiency of mRNA reduction during the RNAi induction of TbMLP1 RNAi cells, quantitative RT-PCR was performed. *T. brucei* MLP1 non induced or tetracycline induced cells (total number of 2.10^7^ cells) were harvested by centrifugation, then RNA was extracted using the RNeasy Mini kit (Qiagen) following the manufactures’ instructions. Collected RNAs were treated with DNase-TURBO (Ambion) for 30 min at 37 °C to digest residual DNA contamination then cDNAs were synthesized using the Super Script III cDNA Synthesis Kit (Invitogen). PCR reaction mixtures were composed according to the manufacturer’s protocol using the LightCycler-480 SYBR Green I master kit (Roche) for a final volume of 15µL. Reactions were carried out in a Roche Real-Time System using the following cycling conditions: 95 °C for 5 min, followed by 44 cycles at 95 °C for 10 s, 58 °C for 10s and 72 °C for 10 s. A reference gene, GPI8, was used to check RNA integrity (see list of primers in S1 Table).

### Immunofluorescence and high-resolution imaging

Transfected cells were harvested and washed with PBS 1X twice, fixed with 4% paraformaldehyde at room temperature. Fixed cells were then washed with PBS 1X and diluted to obtain an appropriate cell density. Cells were adhered to glass slides coated with poly-L-lysine, then neutralized with 100 mM glycine and permeabilized with 0.2% Triton X-100 for 10 min. Slides were blocked 1 h with 1% bovine serum albumin (BSA) in PBS and then incubated with the corresponding primary antibodies diluted in PBS-BSA for 1 hour. Primary antibodies were visualized with the appropriate secondary antibodies conjugated with Alexa Fluor 488, Fluor 546 or ATTO (STED) Secondary Antibody Conjugates (anti-rabbit and anti-mouse secondaries for STED microscopy). DNA was stained with DAPI (Vector laboratories) or Hoechst 33342 (Thermo Scientific). Finally, slides were mounted with Gold® Antifade Reagent (Thermo Fisher-Invitrogen). Cells were viewed by phase contrast, and fluorescence was visualized using appropriate filters on a Zeiss® Axioplan 2 microscope with a 100 X objective. Digital images were captured using a Photometrics CoolSnap CCD camera (Roper Scientific®) and processed with MetaView (Universal Imaging®) or Digital images were captured using an ORCA-flash4.0 camera (Hamamatsu) and processed by ZEN software (Zeiss). Confocal and STED imaging was performed using a quad scanning STED microscope (Expert Line, Abberior Instruments, Germany) equipped with a PlanSuperApo 100 X/1.40 oil immersion objective (Olympus, Japan). For superresulution images, Abberior STAR-RED was excited at 640 nm with a dwell time of 10 µs and STED was performed at 775 nm. Images were collected in line accumulation mode (five lines accumulation). Fluorescence was detected using avalanche photo diodes and spectral detection (650-740 nm). The pinhole was set to 1.0 Airy units and a pixel size of 20 nm was used for all acquisitions.

### Immunoelectron Microscopy

For immunoelectron microscopy GFP-tagged parasites were fixed with 4% paraformaldehyde in phosphate buffer at 4 °C. Cells were then incubated in 0.1% glycine in phosphate buffer, pelleted and embedded in 12% gelatin, cut in small blocks (< 1 mm) and infused 24 h in 2.3 M sucrose on a rotating wheel at 4°C. Gelatin blocks were mounted on specimen pins and frozen in liquid nitrogen. Cryo-sectioning was performed on a Leica UC7 cryo-ultramicrotome, 70 nm cryosections were picked-up in a 1:1 mixture of 2.3 M sucrose and 2% methylcellulose in water and stored at 4 °C. For on-grids immunodetection, grids were floated on PBS 2% gelatin 30 min. at 37 °C to remove methylcellulose/sucrose mixture, then blocked with 1% skin-fish gelatin (SFG, Sigma) in PBS for 5 min. Successive incubation steps were performed on drops as follows : 1) rabbit anti GFP (Abcam 6556) in 1% BSA, 2) Protein A-gold (UMC Utrecht) in PBS 1% BSA. Four 2 min washes in PBS 0.1% BSA were performed between steps. After Protein A, grids were washed four times 2 min. with PBS, fixed 5min in 1% glutaraldehyde in water then washed six times 2 min with distilled water. Grids were then incubated with 2% methylcellulose: 4% uranyl acetate 9:1 15 min on ice in the dark, picked up on a wire loop and air-dried. Observations and image acquisition were performed on a Jeol 1400+ transmission electron microscope on the Electron Microscopy Platform of the University of Montpellier (MEA; http://mea.edu.umontpellier.fr).

### DNA content analysis

To determinate the DNA content in *T. brucei* control non induced cells or TbMLP1 knockdown cells, a propidium iodide (PI) staining method was used. Mid log phase cells were harvested by centrifugation at 10,000 g, for 5 min at 4 °C; 2.10^7^ cells were used for each sample. Cells were washed with PBS 1X, resuspended in 500 µL of ice-cold 70% methanol, vortexed 1 min and incubated at 4 °C maximum 10 days. After centrifugation, cells were resuspended in PBS1X and 0.1 mg. mL^-1^ RNAse A, placed 20 min at 37°C, centrifuged and incubated 10-30 min on ice with 2.5% PI and immediately analyzed on a NAVIOS Flow cytometer (Beckman Coulter). Bicolor flow cytometry to analyze DNA content with S phase cells detection was performed as described elsewhere (50). To detect S phase cells, mid log growing cells were incubated 30 min with 150 μM IdU, collected by centrifugation and washed with 1x PBS. Cells were fixed at −20 °C with a mixture of ethanol and PBS 1X (7:3) and then washed with washing buffer (PBS 1X supplemented with 1% BSA) and DNA denatured with 2 N HCL. To detect incorporated IdU, cells were incubated with anti-BrdU antibody (diluted in washing buffer supplemented with 0.2% Tween-20) for 1h at room temperature then incubated for 1 h with anti-mouse secondary antibody conjugated with Alexa Fluor 488 at room temperature. Finally, cells were resuspended in PBS 1X supplemented with 10 μg. mL^-1^ Propidium Iodide and 10 μg. mL^-1^ RNAse A. Data were acquired on the fluorocytometer "Miltenyi MACS quant” and analyzed with FlowJo software

### FISH and RNA FISH

*T. brucei* cells were fixed in 4% paraformaldehyde, air-dried on microscope immunofluorescence slides and dehydrated in serial ethanol baths (50–100%). Probes were labelled with tetramethyl-rhodamine-5-dUTP (Roche^®^) by using the Nick Translation Mix (Roche^®^). Hybridization was performed with a heat-denatured DNA probe under a sealed rubber frame at 94 °C for 2 min and then overnight at 37 °C. The hybridization solution contained 50% formamide, 10% dextran sulfate, 2 X SSPE, 250 mg. mL^-1^ salmon sperm DNA and 100 ng of labelled double strand DNA probe. After hybridization, slides were sequentially washed in 50% formamide-2 X SSC at 37 °C for 30 min, 2 X SSC at 50 °C for 10 min, 2 X SSC at 60 °C for 10 min, 4 X SSC at room temperature. Slides were finally mounted in Vectashield with DAPI and microscopically examined; more than 200 cells per condition were counted. After inhibition of TbMLP1 by RNAi, the number of copies of chromosome 1 was determined by using DNA probes targeting the alpha- and beta-tubulin genes (51) and chromosome 8 using the M5 ribosomal RNA probe (47). RNA FISH was performed as described in (52). Briefly, cells were harvested and washed once with PBS 1x, placed on a poly-lysine slide to adhere. Adhered cells are then fixed with PFA 4% for 30min at room temperature and then 10 min with 25mM NH4Cl and simultaneously permeabilized and blocked with saponin and BSA. Cells were pre-hybridized for 1 hour with: 2% BSA, 5X Denhartdt’s solution, 4X SSC, 5% dextran sulphate, 35% formamide, 0.5 g.L^-1^ tRNA, 10U/mL RNasin. Followed by overnight incubation with appropriate probe; 2 ng.L^-1^ of Cy3-d(T)20 RNA FISH probe diluted in pre-hybridization solution then washed twice with serial dilutions of SSC solution (4X SSC, 2X SSC and 1X SSC). Nuclei were visualized by adding Hoechst before slides mounting. Cells were viewed by phase contrast, and fluorescence was visualized using appropriate filters on an ORCA-flash4.0 camera (Hamamatsu) and processed by ZEN software (Zeiss).

### DNA combing

Molecular DNA combing and statistical analyses of replication parameters in *T. brucei* Tb427 wild type cells were performed as described previously (53). Asynchronous cell culture was grown to the density between 3-5 x 10^6^ cells/mL and sequentially labelled with two modified nucleosides: 300 μM iodo-deoxyuridine (IdU, Sigma) and 300 μM chloro-deoxyuridine (CldU, Sigma) for 20 min each. Agarose plugs were prepared with a total of 1 x 10^8^ cells resuspended in 100 μL of 1× PBS with 1% low-melting agarose. Each plug was incubated twice in 2 mL of proteinase K buffer (10 mM Tris-Cl, pH 7.0, 100 mM EDTA, 1% N-lauryl-sarcosyl and 2 mg/mL proteinase K) for 24 hours at 45 °C. Agarose plugs were stained with YOYO-1 fluorescent dye (Molecular Probes), resuspended in 100 μl of TE_50_ buffer (10 mM Tris-Cl, pH 7.0, 50 mM EDTA), melted at 65 °C and digested overnight with 10u of β agarase (Sigma Aldrich) at 42 °C. After removal of agarose, the DNA was resuspended in 4 mL of 50 mM MES (2-(N-morpholino) ethanesulfonic acid, pH 5.7) and the DNA fibres were combed as described previously (54) on silanized coverslips prepared by Montpellier DNA combing facility (MDC, IGMM, Biocampus CNRS). Combed DNA and the modified nuclotides were detected with the following combination of the primary antibodies: anti-ssDNA antibody (clone 16-19, Merck Milipore), the mouse anti-IdU antibody (clone B44, Becton Dickinson) and rat anti-CldU antibody (clone BU1/75, Eurobio Scientific). The following secondary antibodies were used: goat anti-rat antibody coupled to Alexa Fluor 488 (Molecular Probes), goat anti-mouse IgG1 coupled to Alexa 546 (Molecular Probes), and goat anti-mouse IgG2a coupled to Alexa 647 (Molecular Probes). The image acquisition was done with a Zeiss Z1 equiped with a ORCA-Flash 4.0 digital CMOS camera (Hamamatsu) and controlled by MetaMorph (Roper Scientific). Statistical analyses of inter-origin distances and velocities of replication forks were performed using Prism 5.0 (GraphPad Software). Statistical significance of the distributions was assessed using the nonparametric Mann–Whitney two-tailed tests that do not assume Gaussian distribution.

## Results

### MLP1 localized in the nuclear basket of NPCs

Subcellular localization of MLP1 in *T. brucei* was assessed by N- or C- terminal epitope tagging. In *T. brucei* procyclic forms (PCFs), TbMLP1 tagged at its N- (Figure 1A) and C-terminal ends (S1Fig. A) was consistently found at the nuclear envelope all along the cell cycle. High resolution STED microscopy further demonstrated the punctuated signal of MLP1 around the nuclear envelope (Figure 1B). We estimated the average diameter of the nuclear basket surrounded by MLP1 molecules ∼ 99 nm (Figure 1C) as compared to ∼120 nm in vertebrates and ∼ 97 nm in yeast (55–57). Immuno electron microscopy of TbMLP1-GFP revealed a nucleocytoplasmic signal near the NPCs which is consistent with a nuclear basket position (Figure 1D). To gain insights into its precise localization, MLP1 was co-expressed with a tagged version of NUP109, a key NUP from the outer ring of the NPC. While the NUP109 localized at the cytoplasmic face of the NPC, MPL1 was found at the inner face of the NE all along the cell cycle (Figure 1E).

**Figure 1:**
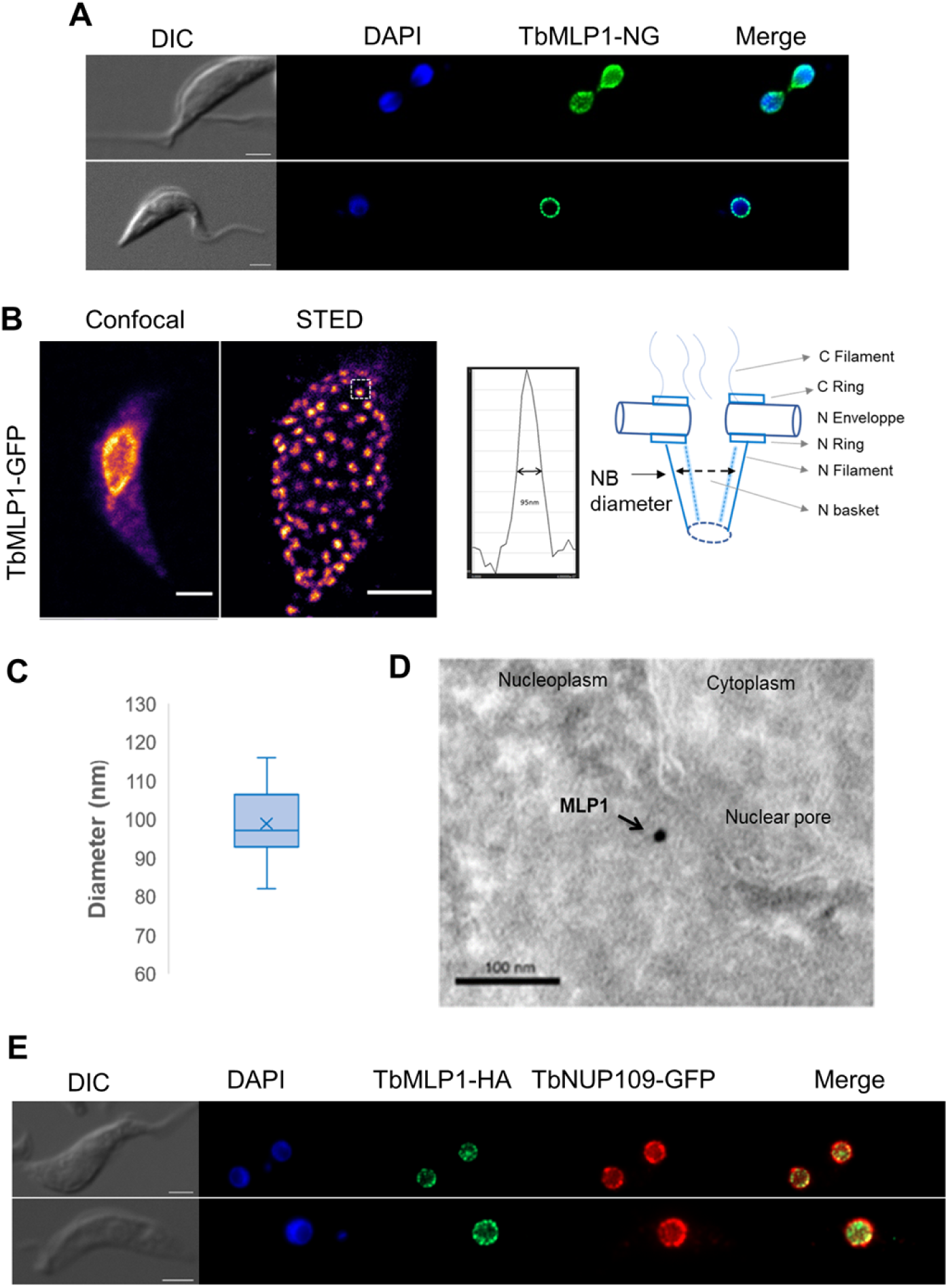
TbMLP1 is a nuclear basket protein localized at the nuclear envelope all along the cell cycle. **(A)** TbMLP1-NeonGreen_n_ localization in *T. brucei* PCFs by immunofluorescence with anti-tag antobodies. DAPI was used to viusalise DNA. Mitotic cells are shown on the upper lane and interphasic cells on the lower lane. Scale bar: 2µm. **(B)** MLP1 localisation was further investigated by 2D superresolution using an ABBERIOR STED superresolution Microscope. The first column shows the standard confocal image and the second the corresponding STED Image. **(C)** Diameter of Pores. Pore diameter measurement was done using FIJI. Data were obtained on >2 slides, >20 nuclei and >100 pores per experiment. **(D)** Immunoelectron microscopy. TbMLP1-GFP was localized in the nucleoplasm beneath a nuclear pore. Gold particle (arrow). Scale bar: 100 nm. **(E)** TbMLP1-NeonGreen_n_ co-expressed with NUP109-HAn in *T. brucei* PCFs. DAPI was used to visualize DNA (blue). Fluorescence was visualized using a Zeiss Z2 microscope and acquired as series of Z-axes. Bar: 2µm for all the presented images.

### MLP1 is essential for cell survival and cell cycle progression

To investigate TbMLP1 functions in *T. brucei* PCFs, we used RNAi-mediated knockdown of MLP1 protein. TbMLP1 was fused with a halotag to follow knockdown efficiency. At day two post induction (D2pi), 48% of cells were TbMLP1-HaloTag negative, 40% had very low labeling and only 12% were TbMLP1-HaloTag positive as compared to 98% TbMLP1-HaloTag positive cells in non-induced (S1 Fig. A-B). IFA results were correlated with qRTPCR results with 50% reduction to TbMLP1 mRNA levels at D2pi (S1 Fig. C). TbMLP1 depletion caused a significant growth defect (Figure 2A) suggesting that TbMLP1 is an essential protein in *T. brucei* PCF cells. To confirm the specificity of the proliferative phenotype we complemented the RNAi strain with either a recodonized HA-tagged version of TbMLP1 (rTbMLP1-HA) or with Lmex.MLP1 from *Leishmania*. The recodonized version rTbMLP1-HA was correctly localized (S1 Fig. D) and growth defect was restored at around 80% (Figure 2A). With the Lmex.MLP1 protein the growth defect was also partially restored (> 50%) (Figure 2A). Next, DAPI staining was used to analyze the NK pattern. In induced TbMLP1 RNAi culture, we observed a severe decrease of typical (1N1K, 1N2K) cells and a significant increase of atypical cells *i.e.* cells with abnormal nucleus (1N*1K: blurred nuclear boundaries and atypical nuclear extensions, enlarged nucleus, micronucleus, nuclear periphery displaying abnormal bulges or blebs) and cells with abnormal dividing kinetoplast morphology and positioning (1N2K* and 2N2K*) (Figure 2B).

**Figure 2:**
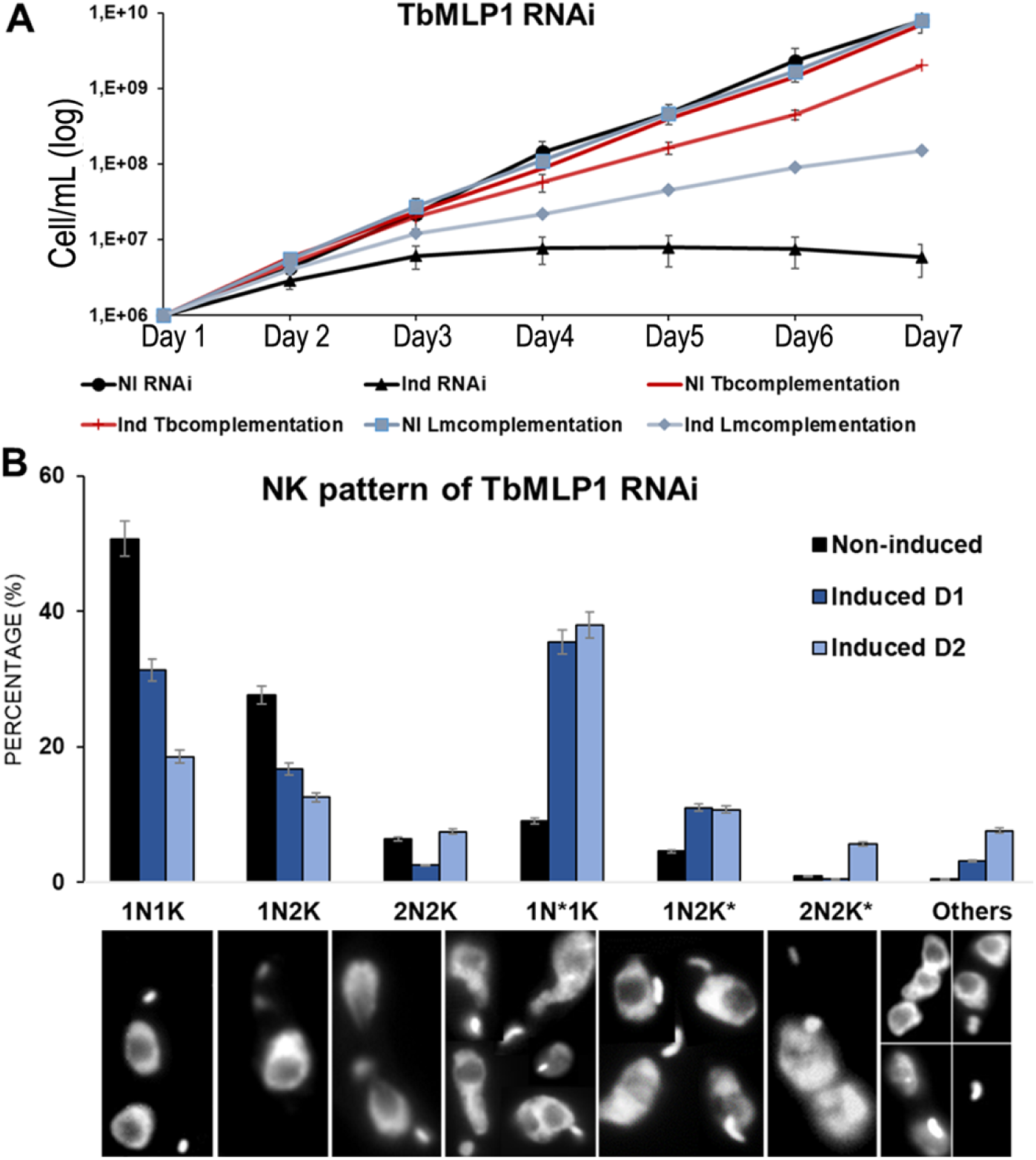
TbMLP1 is essential for parasite survival and correct cell cycle progression. **(A)** Growth curves comparing cell growth of non-induced and tetracycline induced parasites from day one to day six post induction of (i) TbMLP1 RNAi cell line (black lines), (ii) TbMLP1 RNAi complemented cell line with an HA tagged recodonized TbMLP1 version (Tbcomplementation) (Red lines), (iii) TbMLP1 RNAi complemented cell line with Lmex.MLP1 protein (Lmcomplementation) (lightblue lines). **(B)** NK pattern of non-induced TbMLP1 RNAi and induced cell lines. 1N*1K: cells with large, deformed nucleus boundaries and blebbing, 1N2K*: enlarged kinetoplast and nucleus, dividing kinetoplast doesn’t segregate and almost fused with the nucleus, 2N*2K*: cells with defect in segregating the two-daughter nucleus and abnormal kinetoplast positioning.

### MLP1 is required to maintain nuclear envelop integrity but dispensable for NPC architecture

The NK pattern analysis of TbMLP1 depleted cells, allowed us to notice a pronounced increase of abnormal nuclear DAPI staining. We hypothesized that the absence of MLP1 may affect the NPCs and NE architecture. To check for NPCs and NE integrity, we *in situ* tagged NUP109 from the outer ring and NUP98 an FG-NUP. In TbMLP1 RNAi cell line, correct position of NUP109-NeonGreen was observed in 97% of non induced cells, 70% and 35% of induced cells at D2pi and D4pi respectively (Figure 3A and 3C). Indeed, after TbMLP1 induction, atypical cell phenotypes increased with 26% of cells with undetectable NUP109-NeonGreen signal, 30% of cells with DAPI spots surrounded by NUP109-GFP visible in the cytoplasm, at a distance from the nucleus, hereafter called micronuclei, 5% surrounding blebs and 5% of cells with clustered NUP109 and were. Similar results were obtained with NUP98 localization analyses (Figure 3B and 3D). These results suggest that MLP1 is required for accurate positioning of other NUPs and preservation of the NE integrity. The phenotypes observed are therefore reminiscent of those observed after depletion of a trypanosome laminin protein TbNUP-1 which was required for NPC positioning (36). Interestingly, affinity isolation showed that TbMLP1 interacted with TbNUP-1 (38). We hypothesized that TbMLP1 may indirectly affect the NE architecture through an effect on NUP-1 expression or localization. To test this hypothesis, we *in situ* tagged TbNUP-1 in TbMLP1 RNAi cell and followed its expression and localization by IFA. In TbMLP1 non induced cells, TbNUP-1-NeonGreen was correctly localized all along the cell cycle (Figure 4). After tetracycline induction, TbNUP1-NeonGreen localization and targeting to the nuclear envelop was compromised. Thus, at D4pi TbNUP-1 was correctly localized in only 28% of cells. It was mainly present in clusters and some cells totally lost NUP-1-NeonGreen signal, in addition to heterogeneous expression level observed in induced cells.

**Figure 3:**
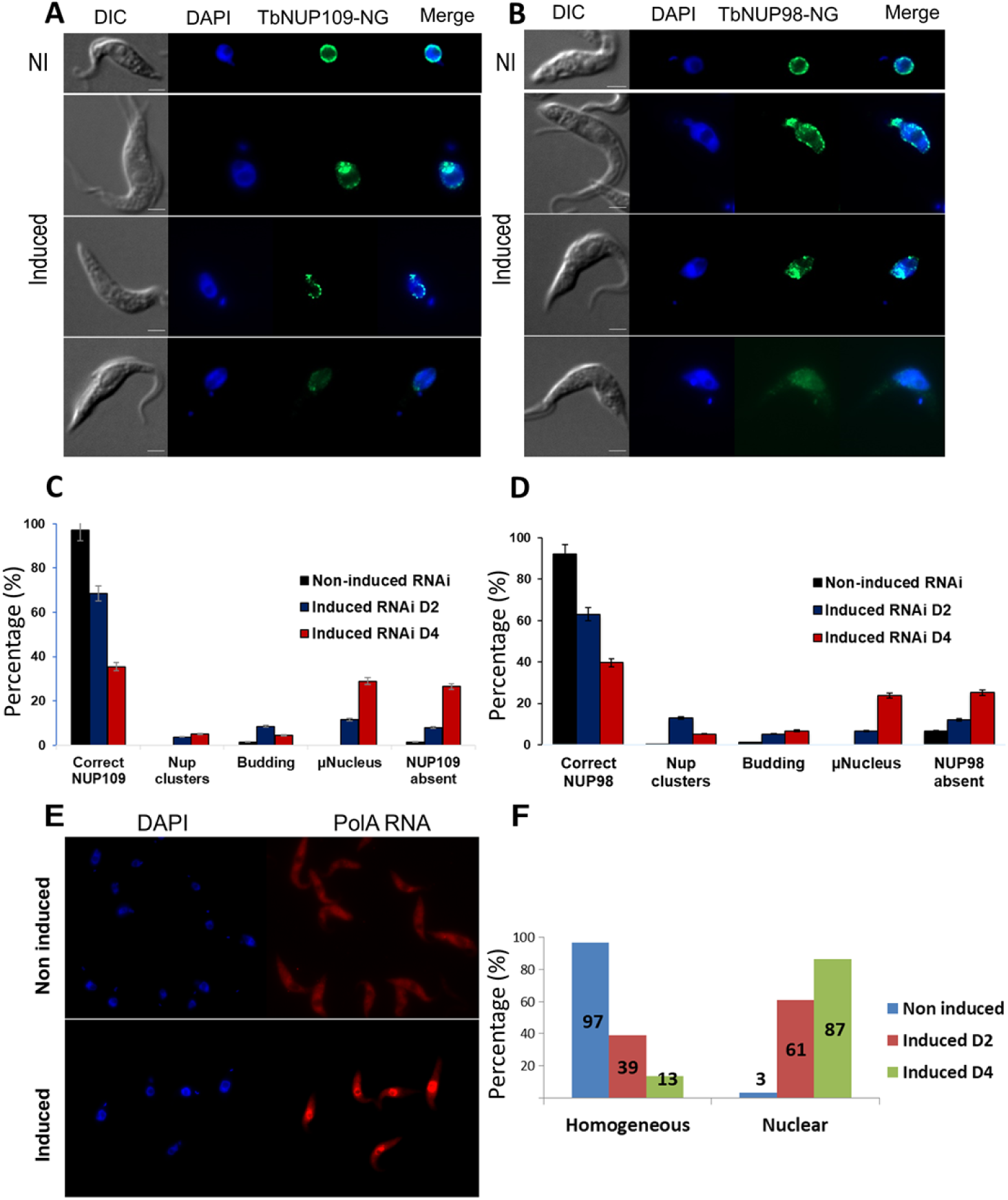
TbMLP1 is required for nuclear pore complex and nuclear envelope integrity. Analysis of TbNUP109-NeonGreen localization pattern all along the cell cycle in non-induced or induced TbMLP1 RNAi cell line (induced D2and D4pi). Upon tetracycline induction a decrease in correct NUP109-NeonGreen localization around the NE and an increase of atypical patterns (NUP clustering blebbing or absence of NUP109-NG labeling and cells with miconuclei). **(A)** Selection of IFA images from TbNUP109-NeonGreen localization in induced TbMLP1 RNAi cells. DAPI was used to visualize DNA (blue). From top to bottom: Induced cells present nuclei with diffuse abnormal extensions or invaginations (blebs), NUP clustering asymmetric nuclear size with micronucleus, cells with NUP109 signal almost undetectable. Fluorescence was visualized using a Zeiss Z2 microscope and acquired as series of Z-axes Bar: 2µm. **(B)** Quantitative data. **(C)** IFA images from NUP98-NG localization (green) in Non-induced and induced TbMLP1 RNAi cells. DAPI was used to visualize DNA (blue). Induced cells present nuclei with diffuse abnormal extension or invaginations (blebs), NUP clustering, assymetric nulear size with micronucleus cells with low signal almost undetectable. Fluorescence was visualized using a Zeiss Z2 microscope and acquired as series of Z-axes Bar: 2µm. **(D)** Graph representing analysis of NUP98-NeonGreen localization pattern through the cell cycle in Non-induced or induced TbMLP1 RNAi cell line (induced D2 and D4), upon tetracycline induction a decrease in correct NUP98-NeonGreen localization around NE) and increase of atypical patterns (NUP clustering blebbing or absence of NUP98-NG labeling and cells with miconucleus). RNA FISH analysis performed on TbMLP1 RNAi cells quantifying the RNA distibution and accumulation in non induced and induced cells at Day 2 and 4 post induction. **(E)** Representative microscopic fields (Non induced top line and induced bottom line). **(F)** Histograms presenting the quantititative data.

**Figure 4:**
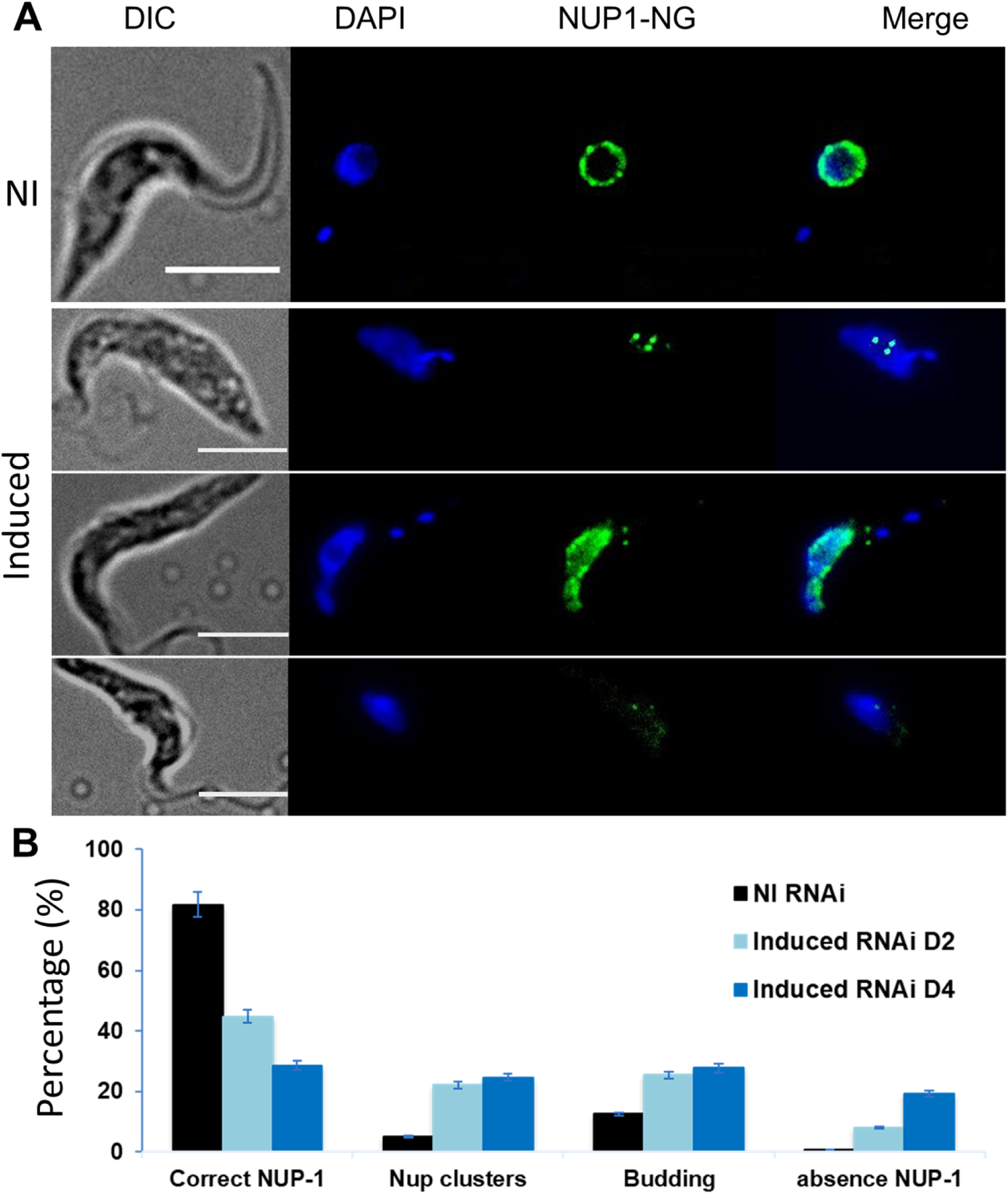
TbMLP1 is required to maintain nuclear envelop integrity. **(A)** IFA images from NUP1-NeonGreen localization (green) in non-induced and induced TbMLP1 RNAi cells. DAPI was used to visualize DNA (blue). NUP1-NG showed punctuated localization around the NE in non-induced cells. Induced cells presented nuclei with diffuse abnormal extensions or invaginations (blebs), NUP clustering, asymmetric nuclear size with micronucleus, cells with NUP-1 signal almost undetectable. Fluorescence was visualized using a Zeiss Z2 microscope and acquired as series of Z-axes Bar: 2µm. **(B)** NUP1-NeonGreen localization pattern all along the cell cycle in non-induced or induced TbMLP1 RNAi cell line (D2 and D4). Upon tetracycline induction a decrease in correct NUP1-NeonGreen localization around the NE and an increase of atypical patterns (NUP clustering blebbing or absence of NUP-1-NG labeling and cells with miconuclei) were observed.

### MLP1 plays an important role in mRNA transport through NPCs

A canonical role described for nuclear basket proteins from other organisms is the regulation of nuclearcytoplasmic transport of RNAs through NPCs. Indeed, in yeast, MLP1 is involved in the retention of unspliced mRNAs in the nucleus *via* a direct interaction with 5’ splice site (58). Still in yeast, MLP1 and MLP2 are also involved in a negative regulatory mechanism of specific genes in response to mRNA export abnormalities (59). To verify TbMLP1 implication in RNA transport we performed RNA FISH experiment using a polyA probe. In non induced TbMLP1 RNAi cell lines, polyA signal was homogenous in around 100% of the labeled cells (Figure 3E); after RNAi induction only 30% of labeled cells retained the homogenous distribution of polyA signal and the remaining cells showed intense nuclear staining corresponding to intranuclear accumulation of mRNAs (Figure 3F).

### MLP1 is required to maintain genome integrity and ploidy stability

Next, we wanted to check if MLP1 displays non canonical functions. Because MLP1 appeared to be important for cell cycle progression, we analyzed DNA content by propidium iodide staining and performed two-color flow cytometry analysis. We observed a reduction of cells in S phase and in G2/M phase and an increase of cells in G0G1 and an increase of cells with abnormal DNA content (>2C) (Figure 5A). To explore the cells with a DNA content >2C, we determined the copy numbers of chromosome 1 and chromosome 8 using FISH analysis. First, we validated these two probes on control cell lines, which appeared to be >90% disomic in *T. brucei*. In TbMLP1 RNAi cell line, aneupmoidy was observed (Figure 5B) and the estimated copy number of chromosome 1 in interphasic cells (1N1K) varied in time; disomic cells decreased down to 28% at D4pi, coupled with an increase of aneuploid cells mainly trisomic (∼30%) and monosomic (16%) (Figure 5C). We observed the same tendency with chromosome 8 probe (Figure 5D). Overall, depletion of TbMLP1 led to mosaic aneuploidy in *T. brucei.* FISH also allowed us to analyse dividing (2N2K) cells and to observe that asymmetrical divisions containing an odd number of homologs were increased in MLP1 depleted parasites (Figure 5E). Asymmetrical divisions containing an even number of homologs is observed when there is a segragtion defect (60). An odd number of homologs in the dividing nuclear may testimony a replication defect preceding a loose segregation. We therefore decided to investigate the role of MLP1 in DNA replication process. We followed DNA replication dynamics after MLP1 depletion by DNA molecular combing technique. Some representative bidirectional replication forks used for this analysis are shown in S2 Fig. A. We compared the replication fork velocities in *T. brucei* before and three days post induction. The analysis showed that there is no statisticaly significant difference in the replication fork velocities (S2 Fig. B). We also performed the analysis of inter-origin distancies (IOD) and this analysis showed that there is no statisticaly significant change in IODs (S2 Fig. C). Our data therefore indicates that TbMLP1 protein is not involved in the regulation of DNA replication, since the dynamics of this process is the same before and after MLP1 depletion.

**Figure 5:**
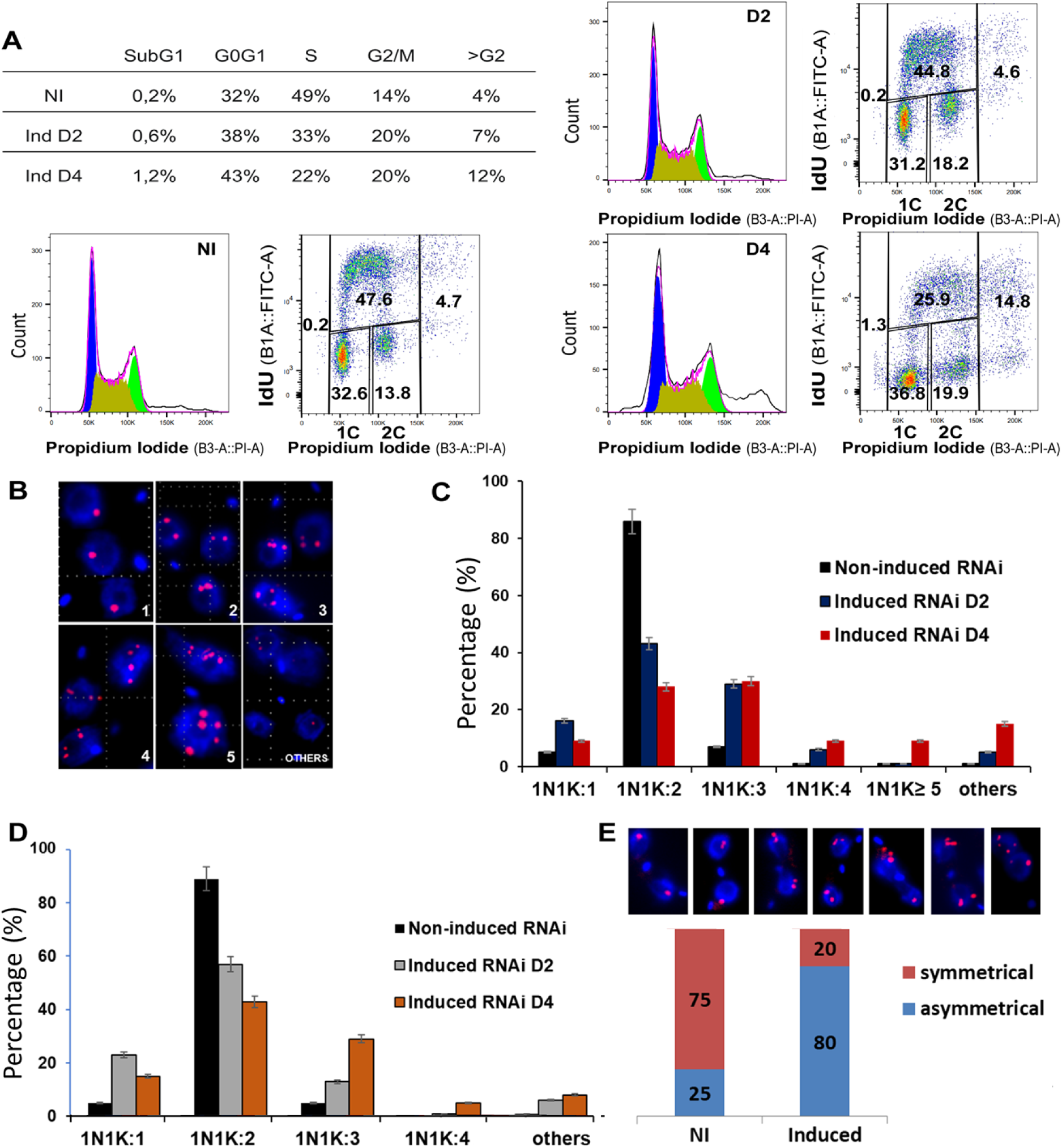
TbMLP1 depletion affects DNA contents and genome stability in T. brucei. **(A)** DNA contents analysis by IP staining (left panels) and two-color flow cytometry (right panels) of TbMLP1 RNAi non-induced cells and induced D2 and D4. The results in the table present the mean percentage of cells in the different cell cycle phases from replicates; representative dot plots from a single expirent are presented. NI and negative control cells were used to draw gates to discriminate between the different cell phases: subG1, G0G1, S, G2/M and >G2. Cells in G2/M have twice as more DNA content (2C) than cells in G1 (1C). Cells in S phase S are IdU positive and have an intermediate DNA content. **(B)** Representatitive FISH images performed on TbMLP1 RNAi cell line using chromosome 1 specific probe. **(C-D)** Quantitative data of FISH analyses, chromosome copy number and patterns were evaluated in intephasic 1N1K cells using chromosone 1 specific probe **(C)** and chromosome 8 specific probe **(D)**. **(E)** FISH analysis on dividing cells. Represetative images top line and quantitative data.

### MLP1 is required for correct chromosome segregation and spindle assembly

The high rate (80%) of asymmetrical division in TbMLP1 depleted cells suggests that TbMLP1 plays a role in chromosome segregation during mitosis. To address this, we assessed the localization of kinetoplastid kinetochore 2 (KKT2) protein during cell division (49). We analyzed the positioning of KKT2 in cells undergoing apparently normal mitosis (1N2K, 2N2K). In normal cells, KKT2 dots align to form the metaphase plate and then move to the cell poles during anaphase and telophase (49). In 80% of non-induced TbMLP1 RNAi cells, TbKKT2-NeonGreen was visible and its localization followed the normal cell cycle kinetics. However, from D2pi to D4pi, the percentage of KKT2-NeonGreen protein correctly positioned decreased from 80% to 40%; KKT2-NeonGreen signal became barely detectable and showed a scattered distribution between the daughter nuclei, suggesting the presence of lagging kinetochores (49) (Figure 6A, 6B). *T. brucei* undergoes a closed mitosis with the mitotic spindle assembled within the nucleus (61). We next investigated the mitotic spindle formation/regulation, since its depletion already affects KKT distribution. To address this, we followed the spindle behavior in MLP1 depleted cells using cellular localization of a Spindle Associated Proteins (SAP) / orphan kinesin named KIN-F (47, 48) (Tb927.3.2020). In non-induced TbMLP1 RNAi cell line, > 80% of dividing cells (1N2K and 2N2K) showed a detectable KIN-F-NeonGreen signal with the expected localization highlighting the mitotic furrow (Figure 6C). At D4pi, 59% of dividing cells displayed either very low KIN-F-NeonGreen signal or no mitotic spindle shape (Figure 6C and 6D). These results suggest that TbMLP1 depletion also affected the proper formation or organization of the mitotic spindle in *T. brucei.* This phenotype is reminiscent of what have been shown for some proteins important for chromosome segregation and spindle formation like RHO-like TbRHP protein (62) and Tousled-like kinase (TLK1) (63).

**Figure 6:**
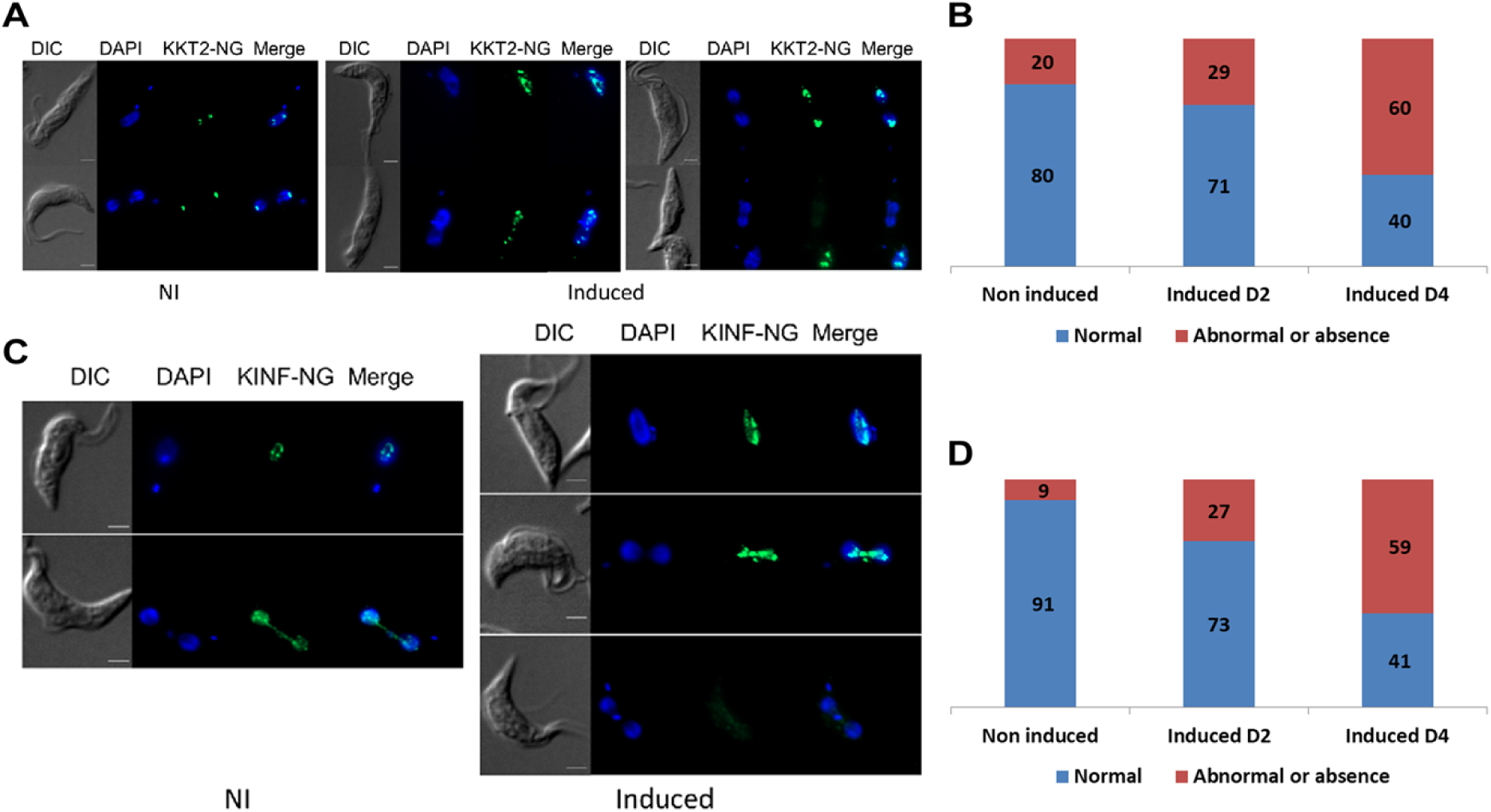
TbMLP1 is required for correct kinetochore distribution during mitosis and for correct mitotic spindle’s network assembly. **(A)** Selection of IFA images from TbKKT2-NeonGreen (KKT-NG) localization in TbMLP1 non induced (NI) and induced RNAi cell line. TbKKT2 is visualized using an anti-NeonGreen anti body (green) and DNA visualized with DAPI (blue). Fluorescence was visualized using a Zeiss Z2 microscope and acquired as series of Z-axes. Bar 2µm. **(B)** Quantitative data of the analysis of TbKKT2-NG localization in dividing cells (1N2K, 2N2K) of NI or induced cell TbMLP1 RNAi cell line (induced D2 and D4). Upon tetracycline induction a decrease in normal TbKKT2 distribution (green) and an increase in abnormal positioning (black) were observed. More than 100 dividing cells were counted for each time point. **(C)** Selection of IFA images from KIN-F-NeonGreen (KIN-F-NG) localization in TbMLP1 non induced RNAi cell line. KINF was visualized using an anti-Neongreen antibody (green) and DNA with DAPI (blue) correct localization was observed following the cell cycle kinetics. **(D)** Quantitative data of the analysis of KIN-F-NG localization in dividing cells (1N2K, 2N2K) of non-induced or induced TbMLP1 RNAi cell line (induced D2 and D4). KIN-F-NG labeling was classified into detectable and normal localization (Green) or indetectable and abnormal localization (Black). KIN-F-NG showed abnormal localization pattern and almost a total absence. More than 100 dividing cells were counted for each time point.

## Discussion

We identified MLP1as a nuclear basket component of the NPC in *Trypanosoma brucei* and demonstrated that its depletion disrupts multiple essential processes. This work combines high-resolution microscopy, immunoelectron imaging, functional genetics, chromosome-level ploidy measurements, and mRNA distribution analyses to build a coherent mechanistic model. The convergence of results from multiple approaches—RNAi, complementation, FISH, flow cytometry, and protein localization—provides strong evidence that TbMLP1 is essential for parasite viability and plays a central role in cell cycle progression. Depletion of TbMLP1 caused accumulation of messenger RNA within the nucleus, nuclear envelope defects, perturbation of kinetochore positioning (KKT2) and mitotic spindle organization (KIN-F) and widespread chromosome segregation errors.

In different models, several NUPs, particularly those forming the nuclear basket, are involved in a wide range of nuclear processes such as nuclear organization and genome integrity (9), cell cycle progression and chromosome segregation ((19–21) and reviewed in (22, 23)). In a previous study, we have shown that TbMLP2 is involved in chromosomal distribution during mitosis in trypanosomatids (39). The phenotypes we observed—aberrant nuclear envelope morphology, micronuclei formation, and chromosome mis-segregation—resemble defects reported in other systems when nuclear basket components are impaired (64–69). Thus, TbMLP1 emerges as a conserved functional counterpart of Tpr/MLPs in opisthokonts (19–21), linking NPC structure to both nuclear architecture and segregation fidelity in kinetoplastids. Although our experiments were performed in the procyclic stage, the fundamental processes affected—mRNA transport, nuclear envelope integrity, kinetochore positioning, and chromosome segregation—are core cellular functions likely conserved across life-cycle stages and potentially in related parasites. Combined with the rescue obtained using *Leishmania* MLP1, our findings suggest that the role of MLP1 is broadly conserved within trypanosomatids despite strong sequence divergence. The essential nature of TbMLP1 and its involvement in multiple core pathways highlight the nuclear basket as a potential vulnerability. While further work is required in bloodstream forms and in infection models, nuclear basket components may represent novel points of therapeutic intervention by targeting nuclear integrity rather than metabolic pathways.

## Acknowledgements

We are deeply indebted to TritrypDB and vEupathDB without which our research would be merely impossible. We acknowledge Marjorie Drac from the Montpellier DNA Combing Facility for providing silanized cover slips. We also acknowledge the MRI platform fo its role in microscopy and flow cytometry acquisitions and analysis. This study was supported by the Agence Nationale de la Recherche within the frame of the “Investissements d’avenir” program (ANR-11-LABX-0024-01 “PARAFRAP”), the Centre National de la Recherche Scientifique (CNRS), the French Ministry of Research and the Centre Hospitalier Universitaire of Montpellier.

## Supporting information captions

### Supporting Information

**S1 Fig.**
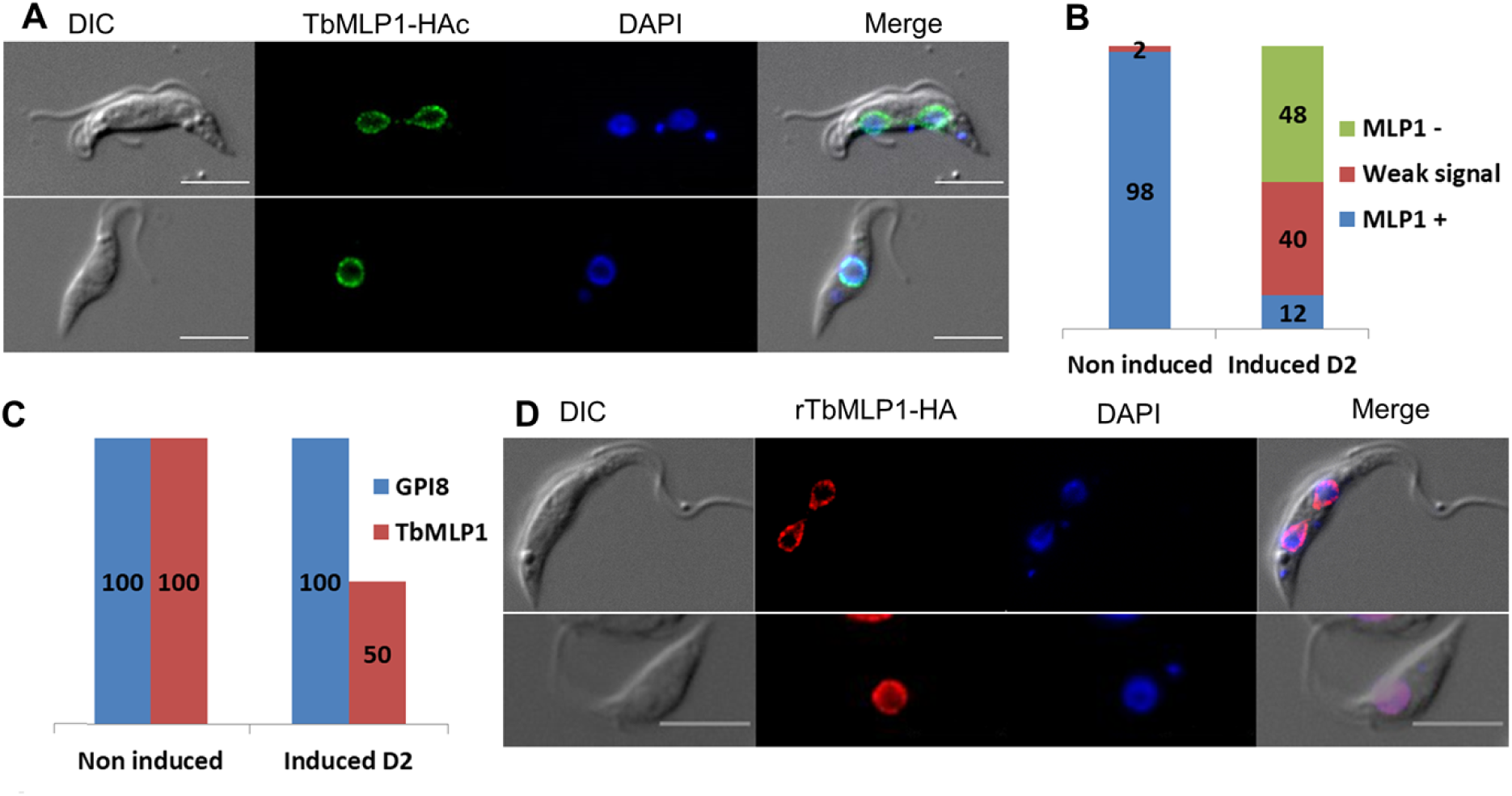
MLP1 sub-localization in T. brucei. **(A)** TbMLP1-HAc localization in *T. brucei* PCF using and anti-HA antibody (Green). DAPI was used to visualize DNA (blue). Mitotic cells (upper lane), interphasic cells (lower lane). Bar: 5 mm. Fluorescence was visualized using a Zeiss Z2 microscope and acquired as series of Z-axes. **(B)** TbMLP1-HaloTag depletion calculated in non-induced RNAi parasites (NI) or two days post tetracycline induction (D2pi) parasites. TbMLP1-HaloTag signal was detected using immunofluorescence assay. TbMLP1-HaloTag labeling was classified into three categories; positive signal (green), weak signal barely detected (gray) and negative signal (blue). **(C)** qRTPCR results performed on cDNAs prepared from TbMLP1 RNAi cell line either non-induced (NI) or induced two days with tetracycline (ind), using primers specific to TbMLP1 and, as a control, primers specific to the housekeeping gene GPI8. **(D)** TbMLP1-HA recodonized version (rTbMLP1-HA) localization in TbMLP1 RNAi cell line visualized with an anti-HA antibody (Red). DAPI was used to visualize DNA (blue). interphasic cells (upper lane), mitotic cells (lower lane).

**S2 Fig.**
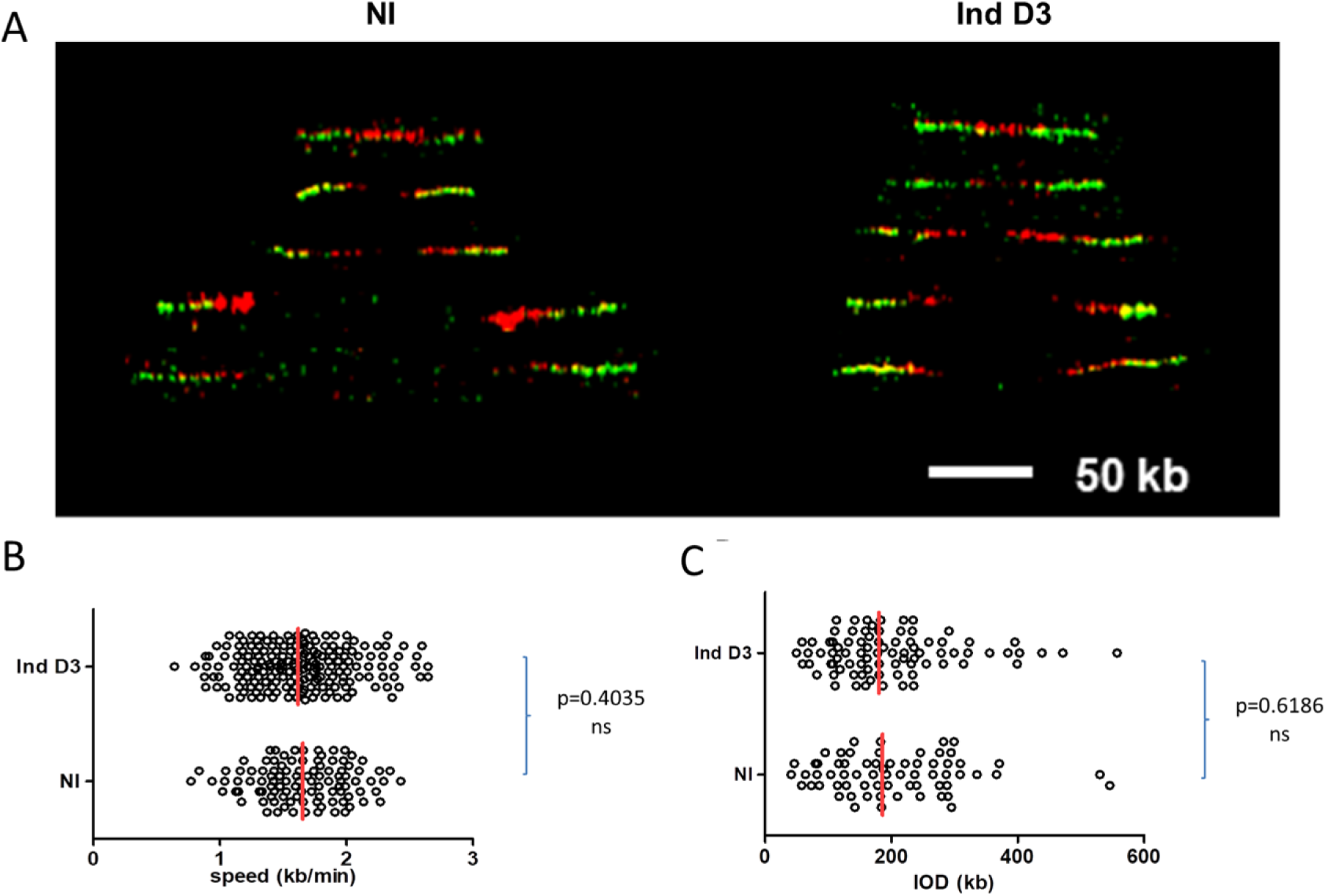
Single-molecule analysis of velocity of replication forks and inter-origin distances (IOD) in TbMLP1 parasites using DNA molecular combing. **(A)** Representative bidirectional replication forks from TbMLP1 RNAi non-induced (NI) or tetracycline induced (Ind D3) taken from different microscopic fields, artificially assembled and centred on the position of the presumed origins. Red tracks: IdU, green tracks: CldU. Scale bar: 50 kb. **(B)** Comparative analysis of the velocity of replication forks in TbMLP1 RNAi non-induced (NI) or tetracycline induced (Ind D3). **(C)** Comparative analysis of the IOD in TbMLP1 RNAi non-induced (NI) or tetracycline induced (Ind D3). p values were calculated using two-tailed Mann-Whitney test (p < 0.05 was taken as significant).

**S1 Table.**
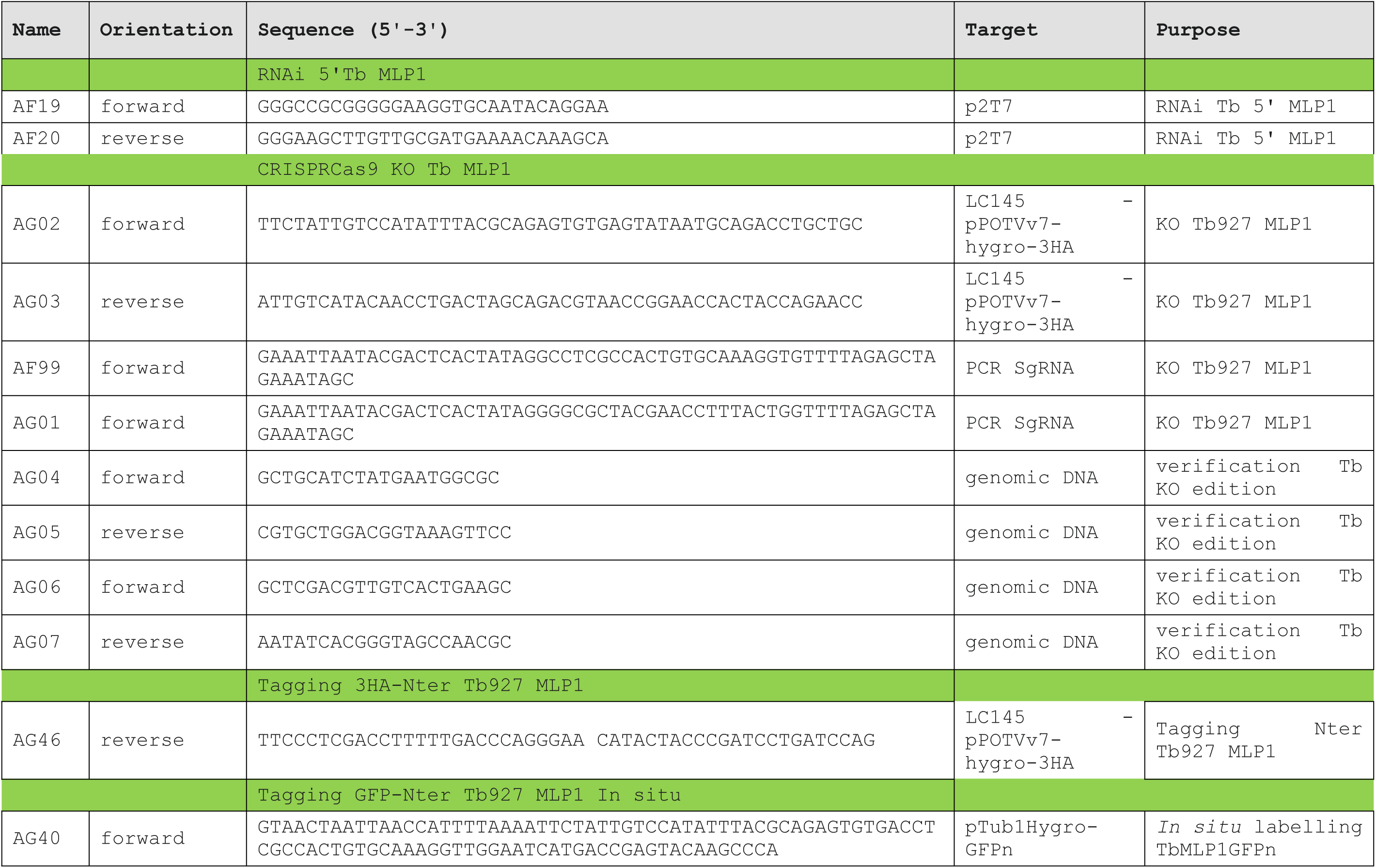
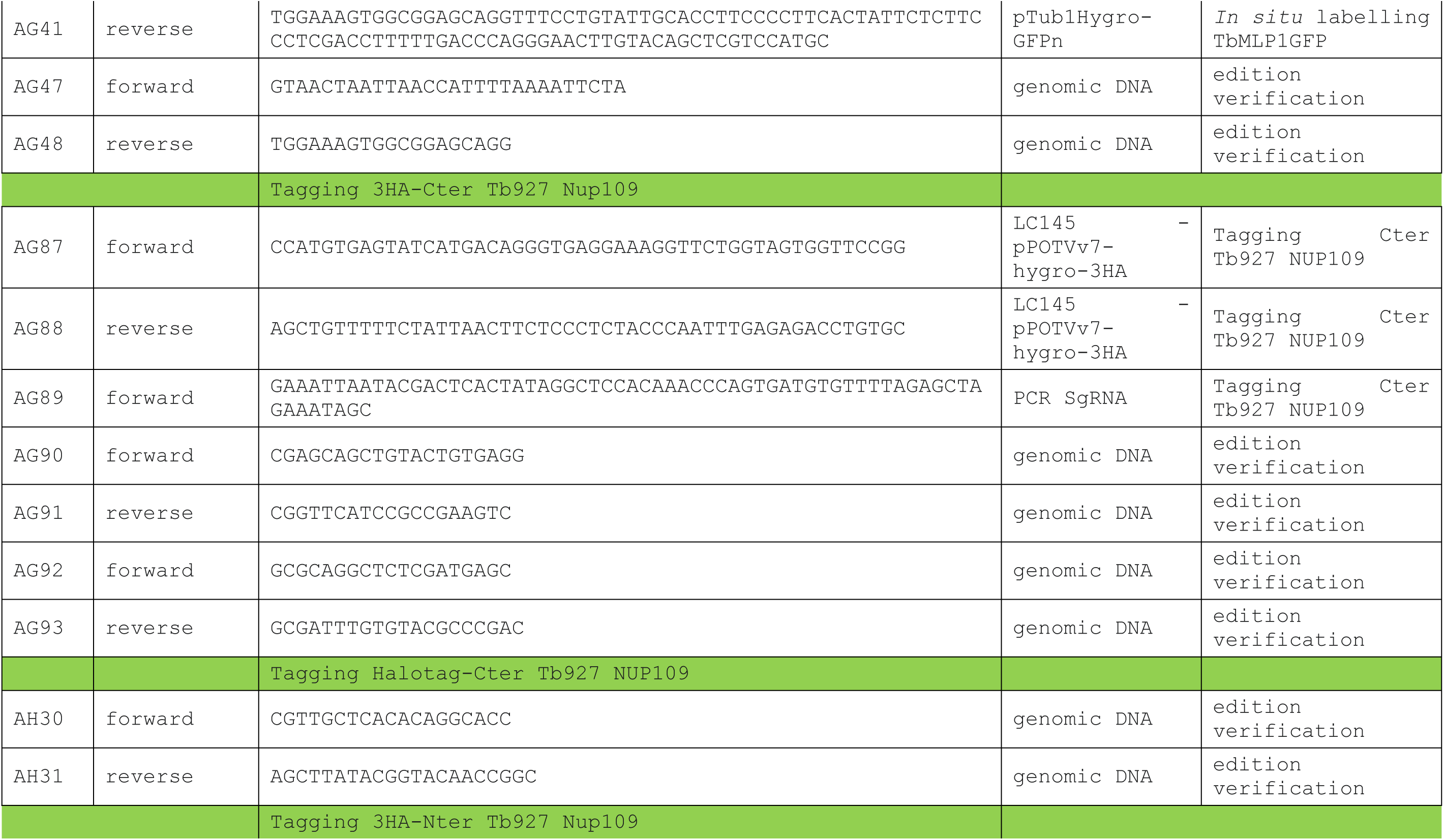
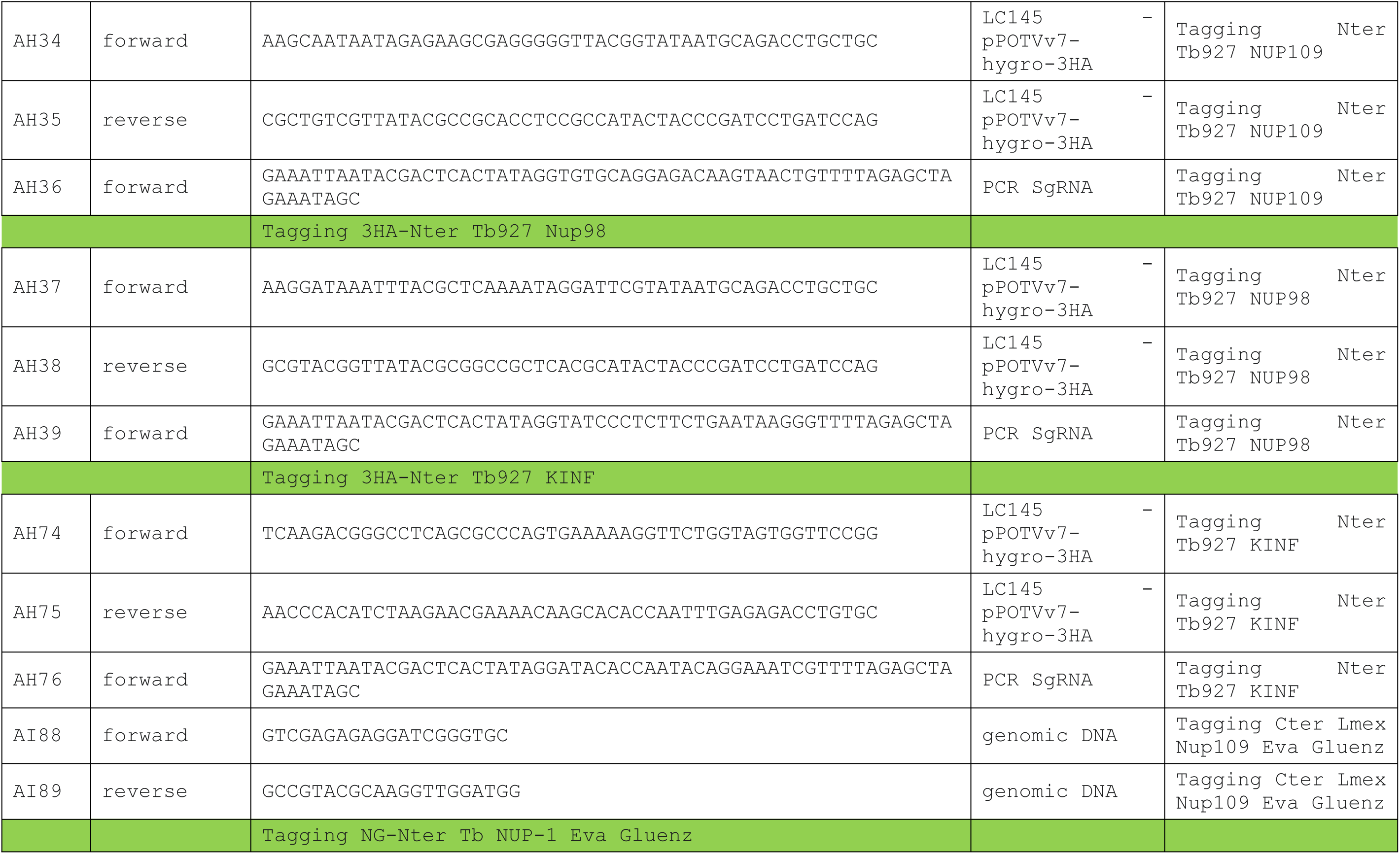
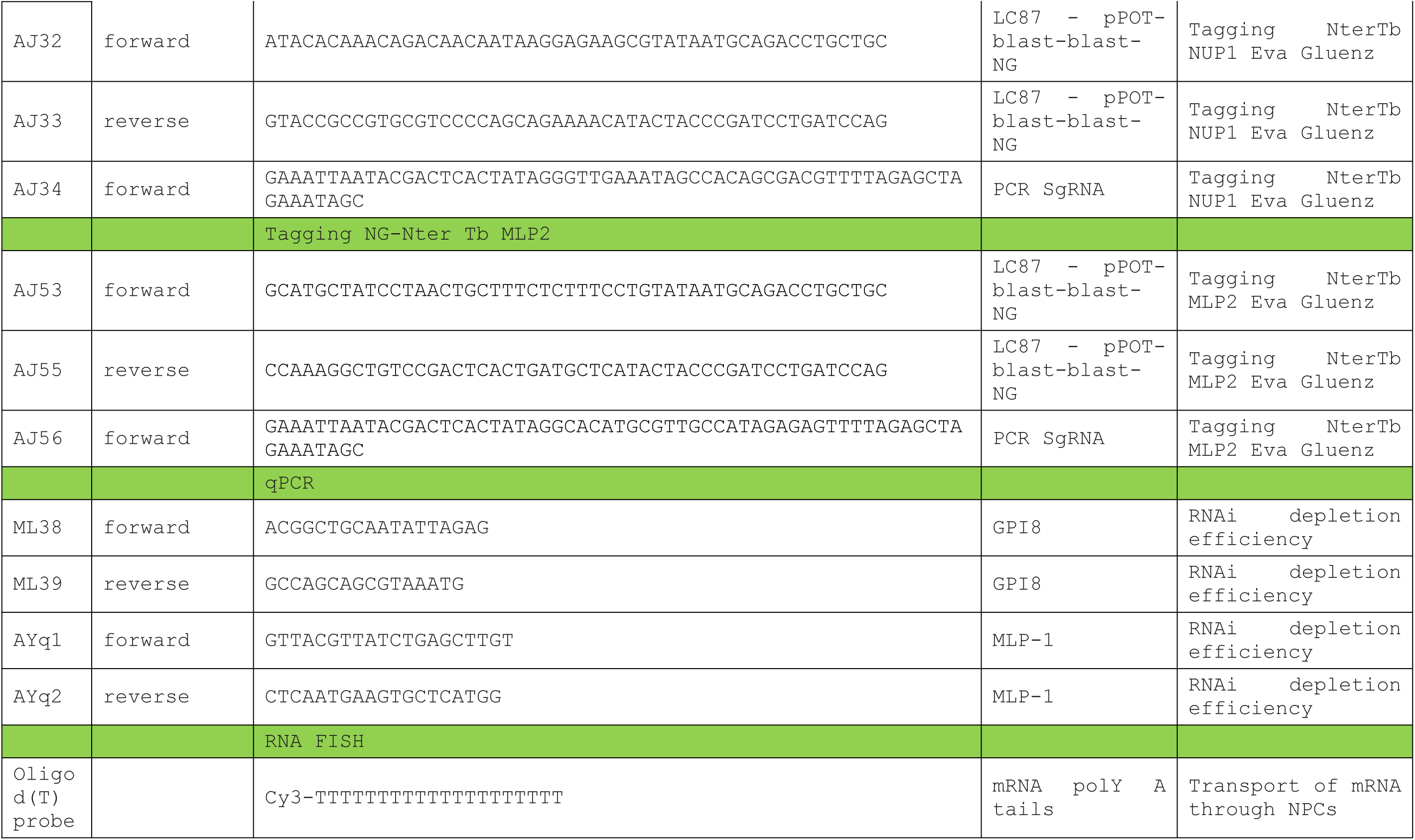
List of primers used for MLP1 characterization.

